# FASTOP - Fast editing toolkit for top expression sites in yeast

**DOI:** 10.64898/2026.05.07.723299

**Authors:** Madhushruti Borah, Nicolas Gautron, Vincent Courdavault, Gita Naseri

## Abstract

Budding yeast *Saccharomyces cerevisiae* is a workhorse chassis for producing added food and agricultural compounds. However, building multi-enzymatic pathways for these chemicals often requires iterative genomic integration, underscoring the need for efficient, rapid genome-editing tools that can reliably target transcriptionally active chromosomal regions. In this study, to accelerate strain construction, we established a genome-editing toolkit to rapidly engineer eight loci, highly expressed hot-spots, but nonessential genomic sites suitable for stable pathway assembly. Our approach integrates three key design features: (i) selectable markers to enable rapid screening of edited cells, (ii) extended homology arms that leverage the yeast homology-directed repair machinery for robust genomic integration, and (iii) co-delivery of Cas9 and guide RNAs to promote efficient double-stranded DNA breaks at specific integration sites. The sequence independence of FASTOP relies on the release of integration cassettes from integrative vectors, mediated by restriction digestion at two flanking multiple-cutting sites in the integration module to minimize the risk of introducing sequence errors during PCR amplification of the integration cassettes. Following the introduction of a fluorescent reporter cassette, we observed high integration efficiencies across the target sites. We then integrated the biosynthetic pathway of plant-derived flavonoid naringenin into the hot-spots of the yeast genome using the FASTOP toolkit. Our results demonstrated that upon expressing the five essential genes in simple shake flask culture, naringenin production reached 505.7 mg/L, representing a significant (69-fold) increase over previously reported titers for comparable minimal heterologous pathways in *S. cerevisiae*. Together, the FATSOP toolkit provides a user-friendly platform for reliably modifying hot-spot loci to rapidly construct multi-enzymatic metabolic pathways in *S. cerevisiae*, while achieving high production levels for high-value food-relevant metabolites.

## Introduction

*Saccharomyces cerevisiae* is an industrial chassis for synthetic biology, which allows the production of value-added compounds and biofuels^1-3^. However, a major limitation for efficient production of heterologous metabolites in microbial hosts is the low transcription of key pathway genes^4-6^, which often results in constrained metabolic flux and decreases strain engineering progress^7^. Overcoming transcriptional bottlenecks remains a central challenge in pathway optimization in a microbial cell factory^4^. The biosynthesis of strictosidine, the central precursor to numerous anticancer natural products in the monoterpenoid indole alkaloid (MIA) family, is a compelling example of this difficulty^5, 8^. Reconstituting strictosidine production in yeast relies on the coordinated expression of a number of heterologous plant enzymes as well as endogenous yeast genes to establish a highly complex plant-derived metabolic network in the microbe^8^. Any imbalance in expression levels can severely limit pathway performance^9^. Notably, the remarkable ~33-fold increase in strictosidine production was achieved by increasing the copy number of genes encoding rate-limiting enzymes^10^. An alternative, potentially more streamlined strategy than introducing multiple gene copies is to integrate pathway genes into genomic loci with intrinsically high transcriptional activity.

Strategic integration into high-expression “hot-spots” in the yeast genome can enhance transcript abundance while maintaining genomic stability. Our understanding of chromosomal positional effects on gene expression in *S. cerevisiae* has recently been significantly improved^11, 12^. This has enabled us to implement new approaches to rational, more predictable strain engineering. While CRISPR/Cas9 has accelerated targeted genome editing, to our knowledge, none of the existing toolkits enable sequence-independent, PCR-free integration into hot-spot genomic sites with high efficiency. In response, we developed FASTOP, a Flexible, Accelerated, Sequence-Independent toolkit for engineering TOP expression sites of the yeast genome. FASTOP implements three key design features: (i) Co-delivery of Cas9 and guide RNAs (gRNAs) to induce site-specific double-strand DNA breaks (DSBs) at any target loci; (ii) Extended homology arms (≥500 bp) to enhance homology-directed repair (HDR) efficiency by native yeast precise DNA repair machinery; and (iii) Yeast selection markers for screening of edited clones, to enable identification of successful integrants without the need for sequencing.

Crucially, FASTOP enables sequence-independent cassette release using restriction enzymes, eliminating the need for PCR amplification and reducing the risk of sequence errors. Eight desirable intergenic loci, located adjacent to the highest reported expressed genes, were fully characterized for three criteria: 1) CRISPR-mediated GFP integration efficiency; 2) expression level, assessed by GFP fluorescence; and 3) multienzyme pathway performance, evaluated through production of naringenin, a flavonoid known for its antioxidant, anti-inflammatory, and applications in functional foods and nutraceuticals^13, 14^, which was more than 69-fold higher than previously reported titers for comparable minimal heterologous pathways. In summary, FASTOP provides a user-friendly, scalable, and high-performance genome-editing platform for the high-level expression of heterologous genes in yeast.

## Materials and Methods

### General

Plasmids were assembled using NEBuilder HiFi DNA assembly (New England Biolabs, Frankfurt am Main, Germany)^15, 16^. All gBlocks, primers, and oligonucleotides were ordered from IDT (Integrated DNA Technologies Inc., Dessau-Rosslau, Germany). PCR amplifications were done using Q5 DNA Polymerase (New England Biolabs, Frankfurt am Main, Germany), or PrimeSTAR GXL DNA Polymerase (Takara Bio, Saint-Germain-en-Laye, France) according to the manufacturer’s recommendations. Amplified DNA parts were gel-purified prior to further use. All constructs were confirmed by sequencing (GENEWIZ, Germany). Primer and plasmid sequences are given in **Supplementary Table S1** and **Supplementary Table S2**, respectively. All strains generated in this study are listed in **Table 1**.

**Table 1.** List of parental *S. cerevisiae* strains used in this study.

| Yeast strain | Relevant genome | Origin |
| --- | --- | --- |
| YPH500 | MAT a, ura3-52, lys2-801, ade2-101, trp1D63, his3D200, leu2D1 | ATCC, #76626 |
| Pro<sub>TEF1</sub>-yEGFP | YPH500/ura3-52::P<sub>TEF1</sub>-yEGFP-T<sub>CYC1</sub>-his3-ura3 | This study |
| yMA001 | YPH500/YBR128C::P<sub>TEF1</sub>-yEGFP-T<sub>CYC1</sub>-leu2 | This study |
| yMA002 | YPH500/YDR448W-1::P<sub>TEF1</sub>-yEGFP-T<sub>CYC1</sub>-trp1 | This study |
| yMA003 | YPH500/YDR448W-2::P<sub>TEF1</sub>-yEGFP-T<sub>CYC1</sub>-trp1 | This study |
| yMA004 | YPH500/YGR240C-1::P<sub>TEF1</sub>-yEGFP-T<sub>CYC1</sub>-lys2 | This study |
| yMA005 | YPH500/YGR240C-2::P<sub>TEF1</sub>-yEGFP-T<sub>CYC1</sub>-lys2 | This study |
| yMA006 | YPH500/YHR142W-1::P<sub>TEF1</sub>-yEGFP-T<sub>CYC1</sub>-natMX | This study |
| yMA007 | YPH500/YHR142W-2::P<sub>TEF1</sub>-yEGFP-T<sub>CYC1</sub>-natMX | This study |
| yMA008 | YPH500/YDR448W::P<sub>PGI1</sub>-PhCHI-T<sub>PGI1</sub>-P<sub>FBA1</sub>-AtC4H::L5::AtATR2-T<sub>FBA1</sub>-trp1 | This study |
| yMA009 | yMA008/YHR142W::P<sub>PGK1</sub>-At4CL-2-T<sub>PGK1</sub>-P<sub>TDH3</sub>-HaCHS-T<sub>TDH3</sub>-natMX | This study |
| yFASTOP-NG | yMA009/YBR128C::P<sub>TDH1</sub>-AtPAL-2-T<sub>TDH1</sub>-leu2 | This study |

*Escherichia coli* NEB DH5α (New England Biolabs) was used for the transformation of plasmids. Strains were cultivated at 37°C in Luria-Bertani (LB) medium with a suitable *E. coli* selection marker. Genetic transformation of plasmids or linearized DNA fragments into competent yeast cells was done using the LiAc/SS carrier DNA/PEG method^17^. To enable selection for transformed cells, all strains were cultivated at 30°C in media rich in yeast extract, peptone, dextrose, and adenine (YPDA) or in suitable synthetic complete (SC) media deficient in one or more amino acids. Then, verification of the transformation was done using colony PCR and/or a microplate reader.

### Design and construction of CRISPR-Cas9/gRNA plasmids

The gRNAs for the loci *YBR128C, YDR448W, YGR240C*, and *YHR142W* were designed using CHOPCHOP v3^18^ and Benchling (www.benchling.com). Based on their efficiency and off-target scores, two gRNAs were chosen for each of the above loci. The gRNAs for *H1, H2, H5*, and *H7*^11^, along with the eight gRNAs for *YBR128C, YDR448W, YGR240C*, and *YHR142W*, were synthesized as oligo primers (primers MA01-MA24), as described in **Supplementary Table S3**. These oligos were annealed and T4 ligated into the BsaI-digested pCRCT backbone, which comprises Cas9, tracrRNA, crRNA, and the URA3 yeast selection marker (Addgene #60621), to construct plasmids pHV001-pHV012. Plasmid pHV002 was modified to pHV013 to change the yeast selection marker to *TRP1* by assembling *TRP1* (primers MA25/MA101, on IDT gBlock) into the pHV002 backbone (primers MA26/MA27). Correct sequence integrity of the constructs was verified by colony PCR and Sanger sequencing.

### Construction of donor plasmids

Donors comprising homologous recombination sequences with unique yeast selectable markers were constructed for each locus. Homologous recombination sequences were based on 500 bp flanking the gRNA sequences beyond the protospacer adjacent motif (PAM). For construction of the donor plasmids pIV001-pIV008, 500 bp of left homologous arm flanking towards the 5’ end of gRNAs (primers MA28/MA43, on genomic DNA of *S. cerevisiae* YPH500 strain), yeast selectable markers *HIS3*/*URA3*/*KanMX*/*hphMX6*/*LEU2*/*TRP1*/*LYS2*/*natMX* (primers MA44-MA59, plasmids from lab collection), MCS introduced in the primers and 500 bp of right homologous arm flanking towards the 3’ end of gRNAs (primers MA60-MA75, on genomic DNA of *S. cerevisiae* YPH500 strain) were cloned into *Bam*HI-digested PUC19 vector, as described in **Supplementary Table S3**.

For the construction of the yeast-enhanced green fluorescent protein (yEGFP)-containing donor plasmids pIV009-pIV012, a cassette harboring *TEF1* promoter, yEGFP, and *CYC1* terminator cassette (primers MA76/MA77, on IDT gBlock) was amplified, digested with *Nco*I*-Bam*HI, and cloned into *Nco*I*-Bgl*II-digested plasmids pIV005-pIV008. The sequences were verified, digested with *Xba*I*-Bam*HI, and transformed into yeast strain YPH500 along with the respective gRNA plasmids to generate strains yMA001-yMA007. The transformants were selected in the respective combinations of yeast selection media.

### Flow cytometry and data analysis

Yeast strains carrying *yEGFP* reporter constructs at specific loci were used to assess yEGFP expression at those loci. The set included wild-type (WT) strains, a strain with yEGFP under the constitutive *TEF1* promoter at *the ura3-52* locus, and strains with yeast yEGFP under the constitutive *TEF1* promoter at *YBR128C, YDR448W, YGR240C, and YHR142W loci*. Initial colonies were cultured in selective SC medium, then inoculated into YPDA medium for primary culture and incubated overnight (~16h) at 30°C with agitation at 200 rpm. Secondary cultures were started at an initial OD_600_ of ~0.1 in YPDA medium and grown for 4–6h at 30°C with 200 rpm. Cells were collected by centrifugation, washed with phosphate-buffered saline (PBS), and analyzed by flow cytometry using a Cytek Aurora CS cytometer to measure yEGFP fluorescence, with a minimum of 10,000 cells per sample. FlowJo was used to calculate the geometric mean of yEGFP fluorescence per cell.

### Design and construction of the Naringenin pathway

For the construction of the Naringenin pathway, the coding DNA sequences (CDSs) for genes *AtC4H:L5:AtATR2, PhCHI, HaCHS, At4CL-2, and AtPAL-2*, codon-optimized for *S. cerevisiae*, were used.

The *PGI1* terminator (primers MA78/MA79, on genomic DNA of *S. cerevisiae* YPH500 strain), *PhCHI* CDS (primers MA80-MA81, IDT gBlock9), *PGI1* terminator (primers MA82/MA83, on genomic DNA of *S. cerevisiae* YPH500 strain),

*FBA1* promoter (primers MA84/MA85, on genomic DNA of *S. cerevisiae* YPH500 strain), *AtC4H:L5:AtATR2* CDS (primers MA86/MA87, on IDT gBlock), *FBA1* terminator (primers MA88/MA89, on genomic DNA of *S. cerevisiae* YPH500 strain) were cloned into *Xho*I-digested yeast vector pIV006 for integration into *YDR448W* locus, harboring yeast selection auxotrophic marker TRP1. The generated plasmid was called pIV013.

The *PGK1* terminator (primers MA90/MA91, on genomic DNA of *S. cerevisiae* YPH500 strain), *At4CL-2* CDS (primers MA92/MA93, on IDT gBlock), *PGK1* promoter (primers MA94/MA95, on genomic DNA of *S. cerevisiae* YPH500 strain),

*TDH3* promoter (primers MA96/MA97, on genomic DNA of *S. cerevisiae* YPH500 strain), HaCHS CDS (primers MA98/MA99, on IDT gBlock), *TDH3* terminator (primers MA100/MA101, on genomic DNA of *S. cerevisiae* YPH500 strain), were cloned into *Xho*I-digested yeast vector pIV008 for integration into *YBR128C* locus, harboring yeast selection marker natMX. The generated plasmid was called pIV014.

The *TDH1* promoter (primers MA102/MA103, on genomic DNA of *S. cerevisiae* YPH500 strain), *AtPAL-2* CDS (primers MA104/MA105, on IDT gBlock), and *TDH1* terminator (primers MA106/MA107, on genomic DNA of *S. cerevisiae* YPH500 strain) were cloned into *Xho*I-digested yeast vector pIV005 for integration into the *YBR128C* locus, harboring yeast selection auxotrophic marker LEU2. The generated plasmid was called pIV015.

Genes were integrated into yeast by transforming *Sal*I/*BamH*I-, *Sbf*I/*BamH*I-, and *Xba*I/*BamH*I-digested donor fragments from pIV013–15, respectively. Integration was facilitated by the plasmids HV007, pHV011, and pHV005, which express Cas9 and locus-specific gRNAs (see **Results** section). Transformants were selected on SD–Ura–Trp or SD– Ura–Leu medium and screened by colony PCR, followed by Sanger sequencing of the PCR amplicons.

### HPLC

Single colonies were inoculated into YPDA media and grown overnight (~16h) at 30°C with 200 rpm. The cultures were inoculated in 10 ml of YPDA media to an OD_600_ of ~0.2 and grown at 30°C for 16h. The final strain yFASTOP-NG was subjected to metabolite extraction from 10 ml of yeast cultures. The cultures were harvested by centrifugation at 1,000 g at −20 °C for 5 min. Cell pellets were washed with 5 ml of pre-chilled 60% (v/v) methanol (−20 °C), then centrifuged at 1,000 g at −20 °C for 5 min. Pellets were resuspended in 1 ml of pre-chilled 80% (v/v) methanol (−80 °C). Cells were transferred to 2 ml bead-beating tubes containing 0.5 mm beads and disrupted using a homogenizer (8,000 rpm; 3 × 30 s cycles with 90 s rest), followed by incubation at −80 °C for 20 min and a second homogenization step (8,000 rpm; 2 × 20 s cycles with 90 s rest). Lysates were collected and centrifuged at 1,000 g at −20 °C for 30 s, and the supernatant was filtered through a 0.22 µm syringe filter, diluted 10 times in methanol, and centrifuged at 10,000 g for 10 min before analysis. For quantification, the extracts were analyzed using an ACQUITY UPLC system (Waters) equipped with a diode array detector (DAD). The system was controlled using Empower 3 software (Waters). Chromatographic separation was performed on an Acquity HSS T3 C18 column (150 × 2.1 mm, 1.8 µm; Waters) at 55 °C with a flow rate of 0.4 mL min^−1^. The mobile phase consisted of water (solvent A) and acetonitrile (solvent B), both containing 0.1 % formic acid. 5 µL of diluted extracts were separated using a 10 min linear gradient from 85 % to 50 % of solvent A. Naringenin was quantified from UV chromatograms at 285 nm using calibration curves generated from commercial naringenin (Sigma-Aldrich) standard.

### Growth assay

Single colonies were inoculated into YNB complete medium (2% glucose + CSM) and grown overnight (~16h) at 30°C with 200 rpm. The cultures were inoculated in 2 ml of YNB medium to an OD_600_ of ~0.2 and grown at 30°C in a microplate reader for the growth assay. OD_600_ was measured for 32h.

## Results

### Design of the FASTOP toolkit

To enable rapid, multiplexed engineering of transcriptionally active loci in *S. cerevisiae*, we developed FASTOP, a modular CRISPR-based platform for integrating up to eight DNA cassettes into predefined hot-spot genomic sites. The system builds on “Y” loci identified by Wu *et al*.; *YBR128C* on Chr II (*ATG14*, Autophagy-specific subunit of phosphatidylinositol 3-kinase complex I), *YDR448W* on chrIV (*ADA1*, Transcription coactivator of the ADA and SAGA transcriptional adaptor/histone acetyltransferase complexes), *YGR240C* on ChrVII (*PFK1*, Alpha subunit of heterooctameric phosphofructokinase), *YHR142W* on ChrVIII (*CHS7*, protein of unknown function)^12^ and expands them with four additional intergenic “H” hot-spot loci: *H1* on ChrXV (between *ADH1* and *YOL085W-A*), *H2* on ChrXII (between *PDC1* and *STU2)*, H5 on ChrIII (between *PGK1* and *ADP1)*, H7 on ChrI (between *CDC19* and *CYC3*), characterized by Baek *et al*.^11^. No more than two of these loci are located on the same chromosome to minimize chromosomal instability.

To engineer the hot-spot loci, FASTOP includes two plasmid series: (i) integration vectors (pIVs) harboring ~500 bp homology arms flanking the insert(s) of interest and a yeast selection marker, but lacking an origin of replication for yeast, enabling direct transformation of linear donor fragments; and (ii) high-copy 2μ helper vectors (HVs) which encode constitutively expressed Cas9 and locus-specific gRNAs to improve the integration efficiency at the site of choice. For each “Y” site, two gRNAs were designed (see Materials and Methods), and for the “H” sites, we used the gRNAs designed by Baek *et al*. into pHV plasmids. Upon co-transformation, Cas9-induced double-strand breaks stimulate homology-directed repair, enabling precise integration of donor cassettes. The plasmid collection includes matched integration/helper pairs for all H- and Y-series loci, and in combination with our previously established pCOM002 and pCOM009 auxotrophic and dominant marker-recycling system^9^ to permit iterative rounds of genome engineering.

### Assessment of the integration efficiency into host spot loci using the FASTOP toolkit

Baek *et al*. (2021) previously characterized four H-series loci for integration efficiency using CRISPR/Cas9 editing methodology^11^. Here, to enable CRISPR-mediated targeting of these Y-series loci, we designed two gRNAs for each site (see **Materials and Methods**) and cloned them into helper plasmids. A GFP expression cassette was cloned into the MCS of the integration vectors. Following co-transformation of linearized donor DNA (upon digestion of integration vectors) and the corresponding helper vector, growing clones on selective media were screened for GFP fluorescence. To determine the integration efficiency, GFP fluorescence of 20 randomly selected colonies per locus was quantified using a plate reader. WT cells were included to determine the basal fluorescence level. We considered colonies with fluorescence signals above the WT threshold to be correct integrants and randomly selected some to confirm integration efficiency. Across tested Y-series loci, integration efficiencies ranged from 75% to 100% (**Fig. 2**), demonstrating robust CRISPR-assisted cassette integration at these highly transcriptionally active genomic sites. However, at the *YGR240C* locus, we obtained only a few colonies using the CRISPR-assisted integration method. Therefore, 20 required colonies were recovered only once the conventional homologous recombination strategy in the absence of CRISPR-mediated genome editing, offered by the FASTOP toolkit, was employed. This data suggests that chromatin accessibility at this locus may limit CRISPR/Cas9 activity, in agreement with previous reports on the effect of chromatin structure, particularly nucleosome occupancy, to influence Cas9 targeting and cleavage efficiency^19, 20^.

**Figure 1.**
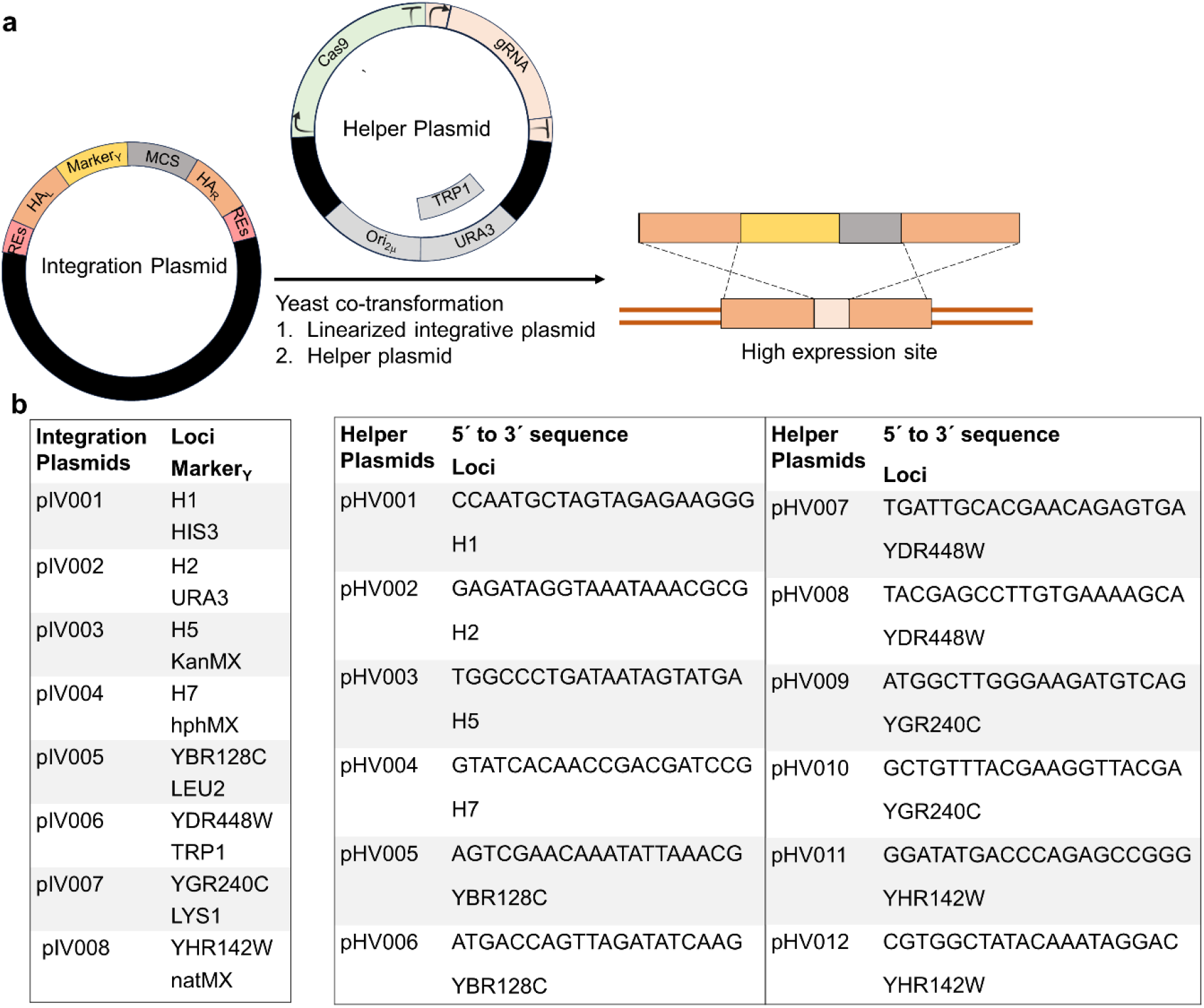
FASTOP toolkit for CRISPR-mediated genomic integration in yeast. **a**Integration vectors contain ~500 bp left and right homology arms flanking a donor cassette with a selectable marker, enabling targeted insertion into predefined chromosomal loci. Episomal helper vectors express locus-specific gRNAs, controlled by the *SNR52* promoter and constitutively expressed Cas9 using the *TEF1* promoter. Following co-transformation of the linearized donor DNA and the corresponding helper vector into yeast cells, targeted integration will occur using a gRNA per locus, and selection markers allow rapid screening of positive integrants. **b** FASTOP plasmid collection. The integration vector series carries donor cassettes with locus-specific homology arms and selectable markers. The helper vector series encodes the matching guide RNA for each hot-spot. Corresponding vector pairs enable modular and multiplex genome engineering. Abbreviations: REs, multiple restriction enzyme sites; L-HA and R-HA, left and right homology arms; MCS, multiple cloning site; Cas9, CRISPR-associated endonuclease 9; gRNA, guide RNA; Ori_2µ_, 2-micron origin of replication; pIV, integration vector; pHV, helper vector; HIS3, URA3, LEU2, LYS1, yeast auxotrophic markers; KanMX, hphMX, natMX, dominant markers.

**Figure 2.**
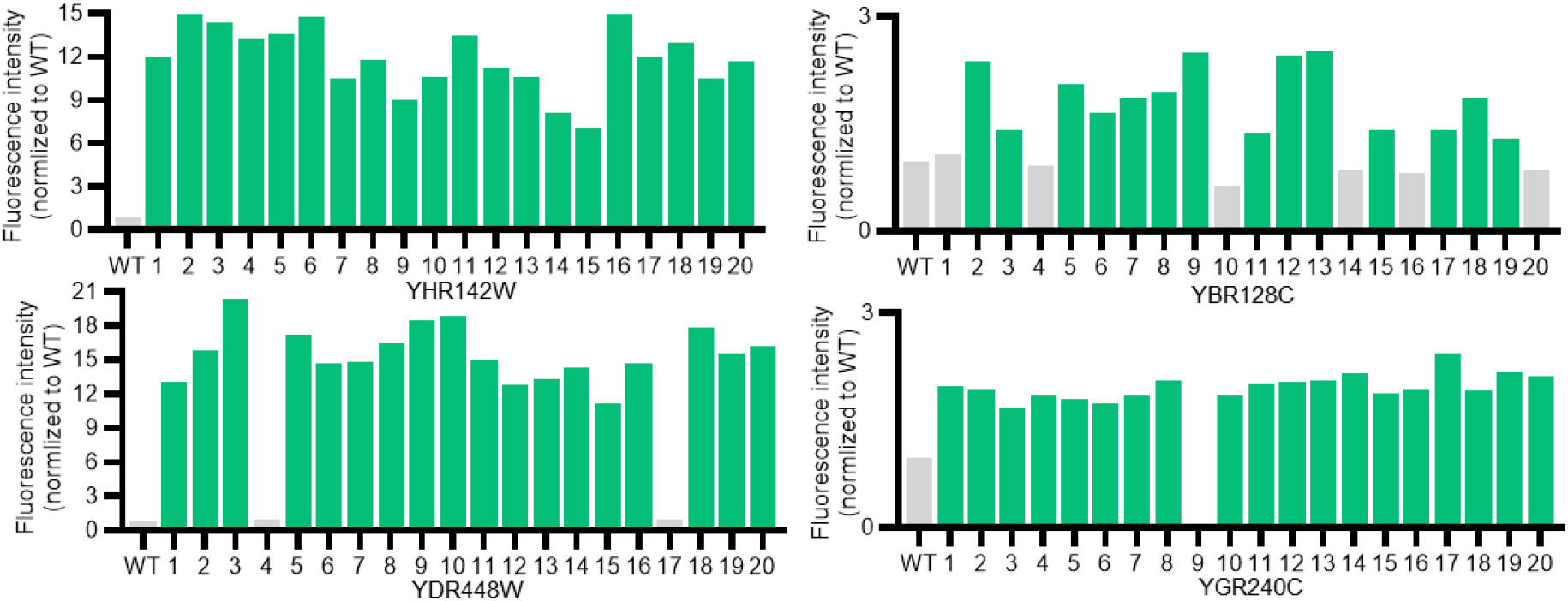
Efficiency of donor integration into hot-spot “Y” loci. To assess the integration efficiency, yEGFP fluorescence intensity was measured for 20 randomly selected colonies per locus using a fluorescence plate reader. *YBR128C, YGR240C, YDR448W*, and *YHR142W* correspond to the Y integration sites^12^. The WT yeast strain was used to determine the basal fluorescence level. Colonies showing fluorescence above the WT baseline were counted as correct integrants. Abbreviations: WT, Wild-type YPH500; yEGFP, yeast-enhanced green fluorescent protein. Full data are shown in **Supplementary Data S1**.

### Characterization of hot-spot loci for gene expression

Baek *et al*. (2021) previously characterized “H” sites for high transcriptional output^11^. Here, to evaluate the transcriptional performance of “Y” genomic hot-spots, the fluorescence intensity of a yEGFP expression cassette integrated into the designated “Y” loci was quantified at the single-cell level and compared with that of the *ura3-52* locus, which served as a previously characterized reference with high expression. Strain YPH500 was used as a negative control to determine background fluorescence levels. As shown in **Fig. 3a**, the “Y” integration sites using CRISPR-mediated genome editing supported strong GFP expression, with *YBR128C, YDR448W*, and *YGR240C* showing fluorescence intensities similar to those observed at *ura3-52*. In contrast, mean fluorescence at *YHR142W* was lower than that of the other “Y” loci and the *ura3-52* reference. Importantly, no significant differences were observed between the two gRNAs tested at each locus (**Supplementary Fig. S1a**), indicating that the observed expression differences were primarily locus-dependent rather than gRNA-dependent. However, single-cell analysis (**Fig. 3b**) revealed greater heterogeneity for the yEGFP cassette integrated into YGR240C via CRISPR genome editing, with approximately 8% GFP-positive cells. Such locus-dependent differences in reporter expression are consistent with previous reports about the chromosomal position effects in *S. cerevisiae*, highlighting that transcriptional output depends on the genomic context of the integration site^21, 22^ and align with our earlier finding (**Fig. 2**) demonstrating that CRISPR-mediated integration was not successful at this locus, due to likely limited Cas9 accessibility to local chromatin. Although relying on homologous recombination for integration into *YGR240C* resulted in significantly lower expression levels than CRISPR-mediated genome editing (**Fig. 3a** and **Supplementary Figure S1b**), it yielded a significantly different result to obtain a hemizygous population (98% yEGFP-positive cells) (see **Fig. 3b** and **Supplementary Figure S1c**).

**Figure 3.**
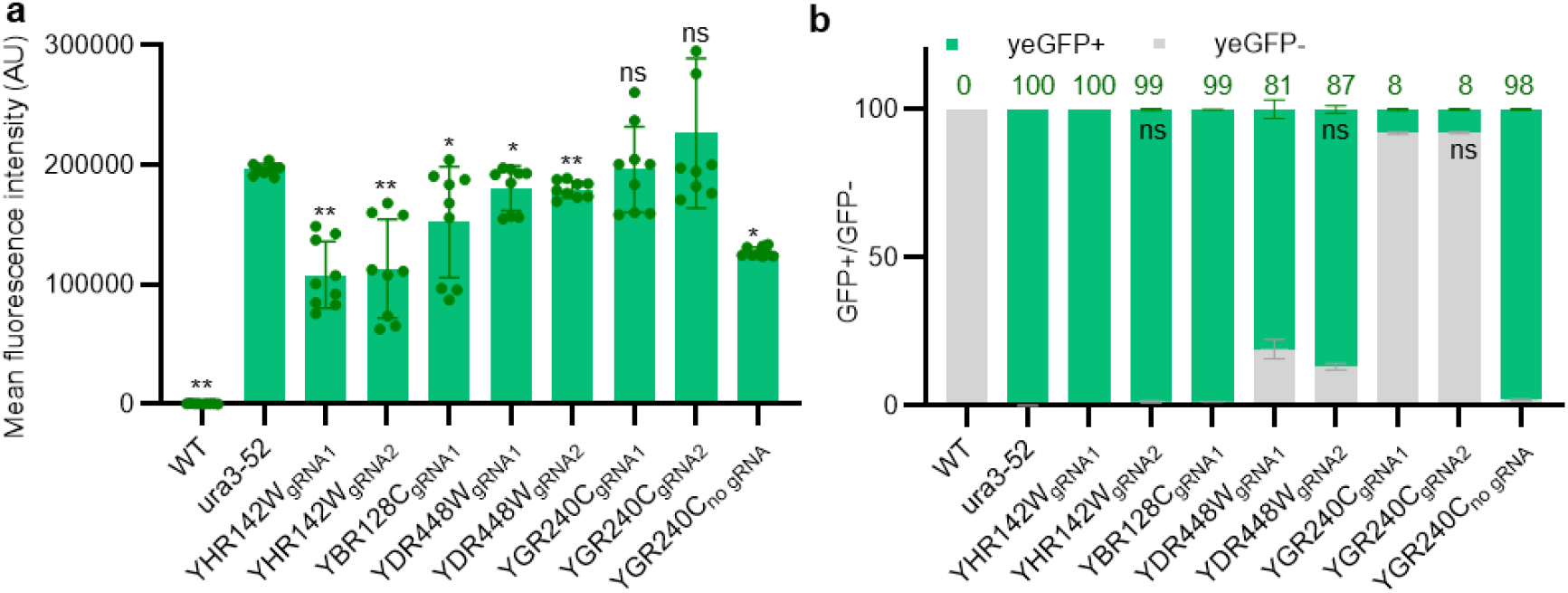
Effect of hot-spot “Y” genomic loci on fluorescent reporter expression in yeast. **a** Transcriptional output of fluorescent protein integrated into the “Y” loci. Cells were transformed with the yEGFP expression cassette for integration into “Y” sites. In the case of CRISPR-based genome editing, for each site, two gRNAs were selected. yEGFP expression in each transformant was measured by flow cytometry, with the *ura3-52* locus selected as a representative of a high-expression site^9^ for comparison. WT YPH500 was used as a negative control. *YBR128C, YGR240C, YDR448W*, and *YHR142W* correspond to the “Y” integration sites^12^. Data represent geometric mean fluorescence intensity ± SD from three biological replicates, each with three technical replicates. Asterisks and ns indicate statistical significance and non-significance relative to the *ura3-52* locus (Student’s t-test; ∗p < 0.05; ∗∗p < 0.01; ns, non-significant). **b** Percentage of GFP-positive to GFP-negative cells determined by flow cytometry. Data represent the mean of the number of yEGFP-positive and yEGFP-negative cells ± SD from three biological replicates, as determined by flow cytometry FACS. Green numbers represent the percentage of GFP-positive cells. ns indicates statistical non-significance relative to GFP-positive cells (Student’s t-test; ∗p < 0.05; ∗∗p < 0.01; ns, non-significant). Abbreviations: FACS, Fluorescence-Activated Cell Sorting; yEGFP, yeast-enhanced green fluorescent protein; gRNA, guide RNA; WT, wild type. Full data are shown in **Supplementary Data S2**.

Baek *et al*. (2021) previously showed that none of the “H” sites selected in this study showed a measurable difference in mean growth rate compared to the WT YPH500 strain^11^. Here, we examined whether the presence of a heterologous DNA in the target “Y” loci could result in any impairment of the growth rate of yeast. The impact of heterologous DNA integration on yeast fitness was examined by measuring the growth rates of each yEGFP-integrated strain in YPDA over 32h, compared to the WT strain and to the commonly used high integration reference site *ura3-52* (**Supplementary Fig. S2**). Integration at *YBR128C* and *YGR240C* resulted in severe growth defects relative to the WT strain, although these strains eventually reached maximum OD_600_ levels comparable to the control. In contrast, integration at the *YDR448W* and *YHR142W* loci resulted in modest alterations in growth dynamics, comparable to those observed at *ura3-52*. Together, our results suggest that the hot-spot sites provide practical genomic integration sites for high expression of heterologous genes in yeast. However, our findings emphasize that selection of genomic integration hot-spots must consider not only transcriptional strength but also influence on cellular fitness.

### Naringenin production in yeast using the FASTOP toolkit

Naringenin is synthesized from phenylalanine and Malonyl-CoA via five enzymatic steps (*AtPAL-2, AtC4H:L5:AtATR2, At4CL-2, HaCHS, and PhCHI*; **Fig. 4a**)^23^. The expression cassettes for *AtC4H:L5:AtATR2* and PhCHI, *HaCHS*, and *At4CL*-2, and *AtPAL-2* were integrated into, respectively, *YDR448W, YHR142W*, and *YBR128C* loci to generate strain yFASTOP-NG (**Fig. 4b**, see also **Method** section). Under simple batch flask culture conditions, the engineered strain produced an naringenin accumulation of 278.24 ± 8.25 mg L^−1^ was obtained (**Fig. 4c**) after 16 hours (mid log-phase, **Supplementary Fig. S2**). Notably, the obtained production level of 505.7 mg/L is 69-fold higher than previous report for comparable minimal heterologous pathways in *S. cerevisiae* yield only 7.3 mg L^−1^ (in the absence of host metabolic rewiring or pathway optimization)^24^. Importantly, despite strong naringenin production, pathway integration at the selected FASTOP loci resulted in slower growth, and biomass ultimately reached a level comparable to that of the WT strain. These results showed that FASTOP-mediated engineering of host-spots can significantly reduce the need for further metabolic engineering strategies such as endogenous pathway rewiring or precursor supply optimization, while still enabling high-level production from a minimal gene set.

**Figure 4.**
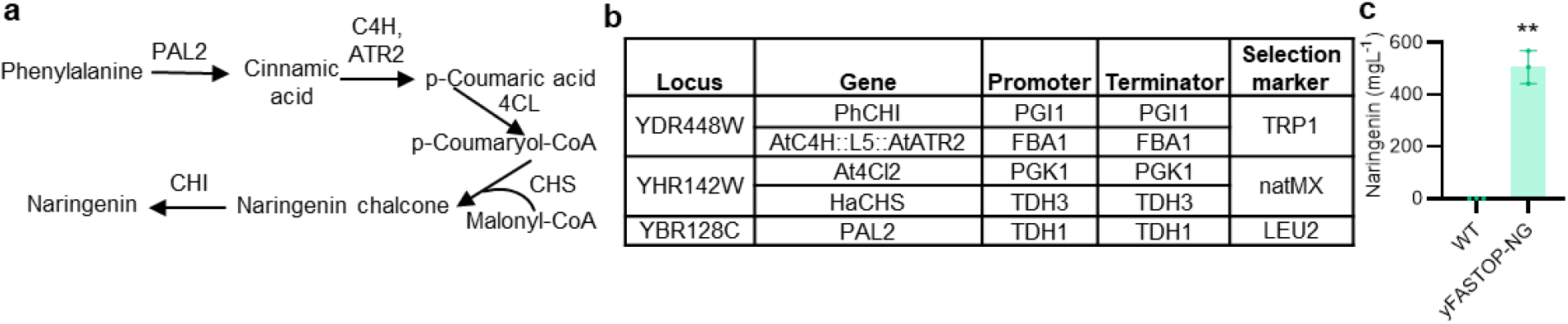
Naringenin production in yeast using the FASTOP toolkit. **a** Schematic illustration of the heterologous naringenin biosynthetic pathway introduced into *Saccharomyces cerevisiae*. Naringenin is synthesized from phenylalanine and malonyl-CoA via five enzymatic steps: AtPAL2, AtC4H-AtATR2, At4CL, HaCHS, and PhCHI. **b** Genomic integration strategy for pathway assembly using FASTOP hot-spot loci. The biosynthetic genes, driven by constitutive promoters, were chromosomally integrated into the *YBR128C, YHR142W*, and *YDR448W* loci to generate strain yFASTOP-NG. **c** Naringenin production in strain yFASTOP-NG. Intracellular naringenin accumulation (mg L^−1^) was quantified at the mid-exponential phase of cells grown in a batch shake-flask. Yeast YPH500 was used as the control WT. Data are presented as mean ± SD from three biological replicates. Asterisks indicate a significant difference from the WT (Student’s t-test; ∗∗p < 0.01). Abbreviations: AtPAL2, phenylalanine ammonia lyase from *Arabidopsis thaliana*; C4H, *Arabidopsis thaliana* cinnamate-4-hydroxylase; ATR2, *Arabidopsis thaliana* cytochrome p450 reductase; 4CL, *Arabidopsis thaliana* 2-coumarate: CoA ligase; HaCHS, *Helianthus annuus* chalcone synthase; PhCHI, *Petunia hybrida* Chalcone isomerase; WT, wild-type. Full datasets are provided in **Supplementary Data S3**.

## Discussion

The development of reliable genome-engineering strategies that enable predictable, high transcriptional output is of high interest for advancing yeast production of food- and agriculture-relevant compounds. This is mainly important as increasing gene copy number, although effective, often introduces instability, recombination risks, and metabolic burden^25^. Hence, integration into naturally active chromosomal environments represents a more stable and streamlined alternative for enhancing pathway expression. Here, we developed FASTOP, a modular CRISPR-based toolkit for rapid engineering of transcriptionally active genomic loci of yeast. The combination of extended homology arms and Cas9-induced DSBs resulted in a high editing success for the tested loci. A notable feature of FASTOP is the elimination of PCR during donor cassette preparation. PCR amplification is widely used in yeast genome engineering, but can introduce point mutations, rearrangements, or sequence bias, particularly when handling long or repetitive constructs, such as those used in engineering projects that rely on synthetic transcription factors and therefore have multiple copies of core binding sites upstream of pathway genes^9, 26^. In these cases, restriction-based cassette release improves sequence fidelity, a feature that becomes particularly important when pathway size and complexity increase. In summary, these features were designed to make the FASTOP toolkit usable in laboratories with varying levels of technical expertise for genome editing in yeast.

While previous genome-wide analyses have revealed strong positional effects on transcription in yeast, only a limited subset of loci has been experimentally characterized for engineering applications. By analytically evaluating the integration efficiency and expression capacity of previously unassessed hot-spot sites, this study provides a practical framework for pathway assembly in the *S. cerevisiae* genome. Here, we validated the four Y-series loci as robust genomic environments for high gene expression. The comparable fluorescence output observed between the “Y” loci and the *ura3-52*, a commonly used reference locus^9^, demonstrates that these regions support high transcriptional activity without requiring multi-copy integrations.

While we have observed a few colonies for YGR240C, when combining both CRISPR/Cas9 and homology-based genomic integration designs, we were able to rescue integration efficiency using only the homology-based approach. This suggests that chromatin accessibility may be an important factor in editing performance, consistent with prior reports that chromatin structure, particularly nucleosome occupancy, can influence Cas9 activity^19, 20^. The toolkit provides a set of plasmids for the engineering of eight host gene genomic sites. Combined with our previously developed marker recycling systems^9^, FASTOP-mediated genomically integrated auxotrophic and dominant selection markers can be efficiently inactivated. As a result, it enables the reuse of FASTOP principles for engineering additional host-spots “H”^11^ and “Y”^12^ for the construction of complex biosynthetic pathways.

Naringenin is a plant-derived flavonoid with well-established antioxidant and anti-inflammatory activities and is used in nutraceuticals and dietary supplements^13, 14^. The successful reconstruction of the naringenin biosynthetic pathway, integrated into three host-spots, resulted in 505.7 mg/L naringenin production, representing a significant (69-fold) increase over previously reported titers for comparable minimal heterologous pathways in *S. cerevisiae*. This data indicates that transcriptionally active genomic environments can support high-level pathway expression without imposing severe fitness costs, suggesting that hot-spot genome integration can potentially reduce the need for extensive host metabolic rewiring or precursor engineering. Distributing pathway modules across separate loci may also reduce transcriptional interference between regulatory elements.

Despite several advantages, to further advance the toolkit, several aspects should be considered, including characterizing the behavior of hot-spot loci under diverse cultivation conditions, characterizing the transcriptional strength of various promoters and terminators integrated into hot-spot loci, and expanding the number of hot-spot loci.

In summary, FASTOP establishes a practical platform for targeted integration into transcriptionally active regions of the yeast genome. It provides a valuable synthetic biology tool for metabolic engineering applications in the sustainable production of food, nutraceutical, and agriculturally relevant compounds.

## Acknowledgements

Gita Naseri acknowledges funding by the DFG (German Research Foundation) Emmy Noether Programm—NA 1650/4-1.

## Author contributions

GN supervised the project. GN conceived the project with contributions from MB. MB designed and performed the experiments. MB and GN analyzed the data. Chromatography experiments were performed and analyzed by VC and NG. The manuscript was written by GN. The Materials and Methods was written by MB. All authors take full responsibility for the content of the paper.

## Data availability

The authors declare that the data supporting the findings of this study are available within the paper and its Supplementary Information files. Should any raw data files be needed in another format, they are available from the corresponding author upon reasonable request. The FASTOP plasmids reported in this study are available from Addgene (www.addgene.org) under the following ID numbers. Integration plasmids: pIV001, #256762; pIV002, #256763; pIV003, #256764; pIV004, #256765; pIV005, #256766; pIV006, #256767; pIV007, #256768; pIV008, #256769; and helper plasmid: pHV007, #256770.

## Supplementary Figures

**Supplementary Figure S1.**
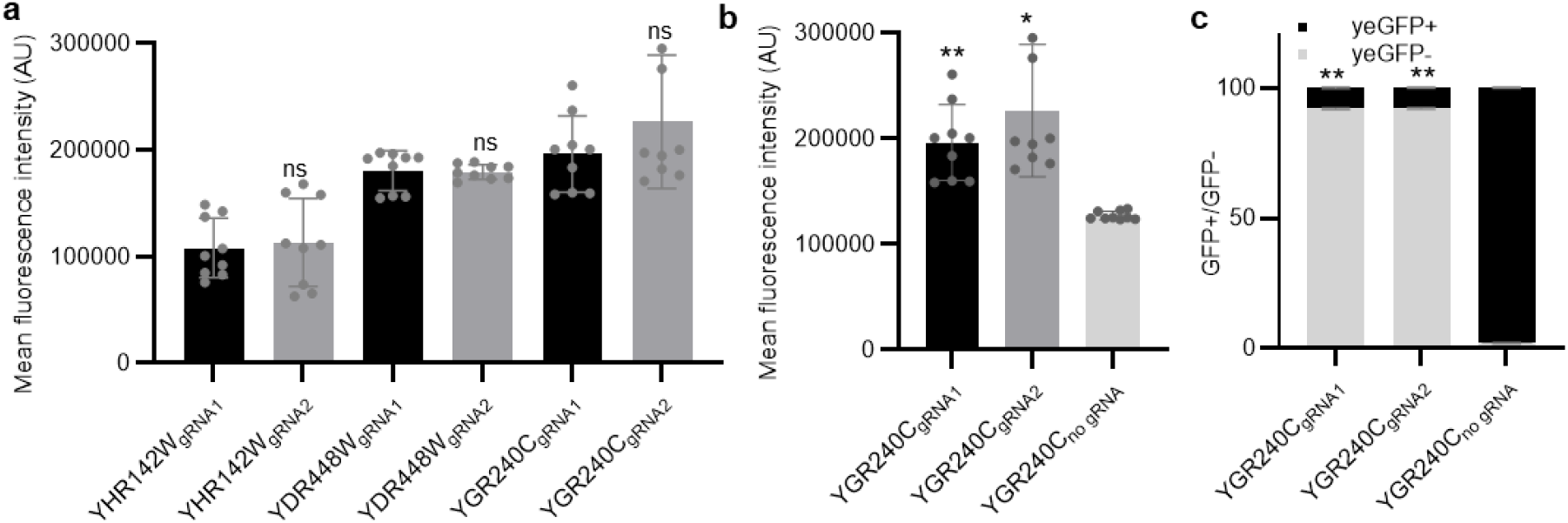
Transcriptional output difference for CRISPR-assisted integration into the “Y” loci using different gRNAs. Cells were transformed with the yEGFP expression cassette to be integrated into “Y” sites. For each site, two gRNAs were selected. yEGFP expression was next measured by flow cytometry FACS. *YGR240C, YDR448W*, and *YHR142W* correspond to the “Y” integration sites^12^. Data represent geometric mean fluorescence intensity ± SD from three biological replicates, each with three technical replicates. **a** There was no non-significant difference between two gRNAs (Student’s t-test; ns, non-significant). **b** Asterisks indicate a significant difference for yEGFP expression level from the condition without gRNA. **c** Asterisks indicate a significant difference for yEGFP-positive cells from the condition without gRNA (Student’s t-test; ∗p < 0.05; ∗∗p < 0.01; ns, non-significant). Abbreviations: FACS, Fluorescence-Activated Cell Sorting; yEGFP, yeast-enhanced green fluorescent protein; gRNA, guide RNA; WT, wild type. Full data are shown in **Supplementary Data S4**.

**Supplementary Figure S2.**
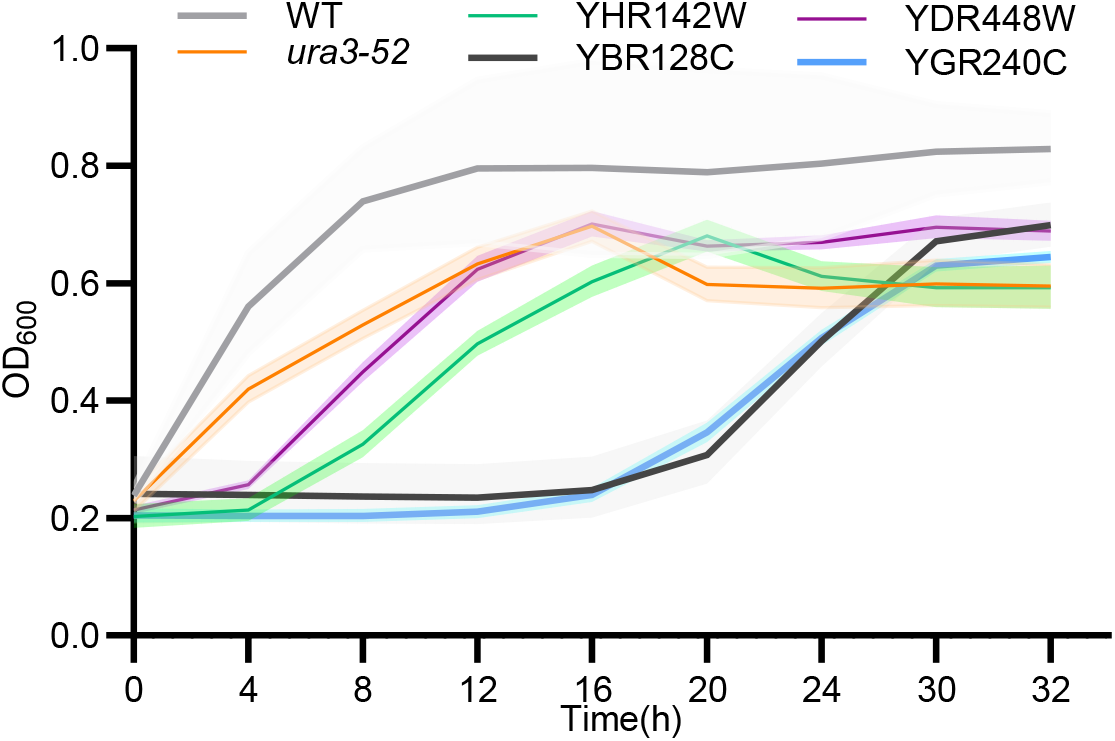
Growth kinetics of yeast strains carrying yEGFP reporters integrated at genomic hot-spot loci. OD_600_ was measured over 32h in YPDA for WT and yEGFP-integrated hot-spot loci strains.*YHR142W, YBR128C, YDR448W*, and *YGR240C* are hot-spot “Y” loci. As controls, WT and strain harboring the yEGFP cassette integrated into the *ura3-52* site were used. Measurements were obtained from three independent colonies per strain and are shown as mean ± SD. Abbreviations: yEGFP, yeast enhanced GFP; OD_600_, optical intensity at 600 nm; h, hour. Full data are shown in **Supplementary Data S5**.

**Supplementary Figure S3.**
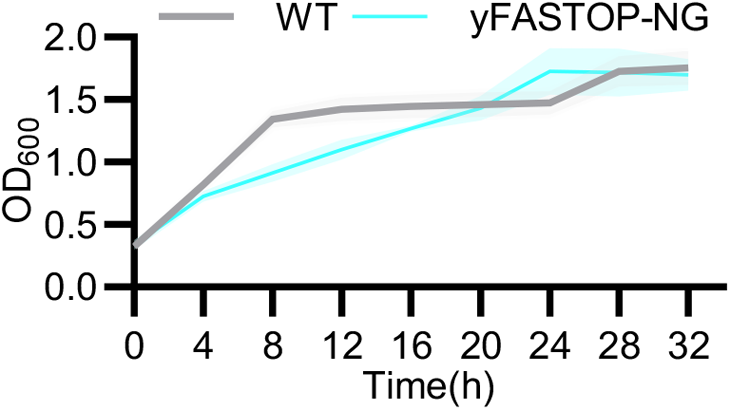
Time-course analysis of the growth of the naringenin-producing strain. The engineered strain was cultivated under batch conditions, and OD_600_ was monitored over time. Yeast YPH500 was used as the control WT. Data are presented as mean ± SD from three biological replicates. Asterisks indicate a significant difference from the WT (Student’s t-test; ∗p < 0.05; ∗∗p < 0.01; ns, non-significant). Abbreviations: WT, wild-type; OD_600_, optical intensity at 600 nm; h, hour. Full datasets are provided in **Supplementary Data S6**.

